# The bully phage: A Shiga toxin-encoding prophage interferes with the induction of co-hosted prophages

**DOI:** 10.1101/2025.01.01.630987

**Authors:** Rachel Hashuel, Tamara Wellins, Maayan Rachimi, Eran Gutman, Reut Melki, Linoy Taktuk, Pe’era Pash Wasserzug, Rinat Avissar, Karina Roitman, Yael Litvak

**Author notes:** These authors contributed equally to this work (Rachel Hashuel, Tamara Wellins).

## Abstract

Shiga toxin (Stx) prophages lysogenize various bacterial species, converting them into dangerous pathogens and driving the ongoing emergence of new outbreaks globally. These pathogens are polylysogens, harboring multiple prophages within their genomes. Yet, the impact of Stx prophage acquisition on resident prophages remains largely unknown. Using *in vivo* transmission studies with the murine pathogen *Citrobacter rodentium* that carries an Stx prophage, we demonstrate that the Stx prophage readily lysogenizes commensal *Escherichia coli* strains within the intestine. Markedly, this lysogenization altered the induction activity of prophages encoded within each new toxigenic host strain. Moreover, the Stx prophage abolished the induction of ϕNP, the most active native prophage of *C. rodentium*. We further show that ϕNP inhibition is mediated by the Stx Cro repressor, which exhibits dual functionality, promoting its own lytic cycle while repressing the expression of ϕNP lytic genes. Our findings reveal a novel competition strategy among prophages, and suggest that interfering with the induction of co-hosted prophages likely plays a role in the emergence and evolution of new Stx-producing pathogens.

## Introduction

Shiga toxin (Stx)-producing pathogens are a significant human health risk, as infections result in hemorrhagic colitis and may lead to kidney failure and death (*1*). The virulence-factor responsible for this severe outcome is the Stx, a highly potent cytotoxin encoded within a lambdoid prophage, a temperate bacteriophage integrated into the bacterial chromosome (*2*–*5*). Prophages are typically repressed to maintain lysogeny, the dormant stage of the viral life cycle. Environmental stressors can trigger prophage induction - a process where the virus switches from lysogeny to the lytic life cycle, leading to active viral replication and production of infective virions. Notably, induction of Stx prophages was shown to be markedly enhanced in the intestinal environment and required for renal disease and lethality (*6, 7*).

Lambdoid prophages remain dormant until the bacterial host encounters stress, such as DNA damage, which activates the bacterial SOS response, an inducible DNA repair system. The SOS response facilitates bacterial survival, but also triggers prophage induction. During lysogeny, the CI phage repressor binds to specific operator sequences within the prophage control region, blocking the expression of lytic genes while promoting its own transcription to maintain the lysogenic state. Following SOS response activation, the bacterial RecA protein becomes activated and facilitates the autoproteolytic cleavage of the CI repressor. As CI levels drop, this relieves repression of the phage-encoded *cro* gene, allowing Cro protein to accumulate. Cro then binds to the operator sites in a pattern opposite to CI, repressing lysogenic genes while allowing expression of lytic genes. The shift in regulatory dominance from CI to Cro initiates the expression of genes required for phage replication and host cell lysis, committing the phage to the lytic pathway and preventing return to lysogeny. In Stx prophages, induction results in lysis of the carrying bacteria and release of both virions and Stx molecules to the surrounding environment (*8*–*11*). The toxin disrupts protein synthesis in epithelial and endothelial cells, causing cell death and epithelial damage that enables it to enter the bloodstream and consequently harm the kidneys and other organs (*12*). The released virions can infect and lysogenize naive intestinal bacteria, facilitating horizontal transfer of virulence genes, thus promoting the emergence of new Stx-producing pathogens. Indeed, phage transmission gave rise to various Stx-producing bacterial pathogens, among them enterohemorrhagic *Escherichia coli* (EHEC), Shiga toxigenic *E. coli* (STEC), Shiga toxigenic *Shigella* species as well as Stx-producing variants of *Citrobacter, Enterobacter*, and *Aeromonas* (*3*–*5, 13*–*18*). Importantly, these pathogens are often polylysogens, carrying multiple prophages, that may constitute up to 13% of their genomes (*19*–*21*). Notable examples include EHEC O157:H7 Sakai, which harbors 18 prophages and was responsible for an outbreak in Japan, and *E. coli* O104:H4, which contains 9 prophages and caused a large outbreak in Europe (*20, 21*).

Upon induction, phages hijack host cell machinery, including transcription and translation systems, as well as essential resources such as nucleotides, amino acids, and energy, to facilitate virion production. This creates competition within a polylysogen host among the encoded prophages. Indeed, not all prophages carried by a bacterial cell replicate following an inducing trigger, and typically, only a subset successfully produce complete virions (*20, 22*). Despite the high transmissibility and recurring lysogenization of new host cells by Stx phages, and although the resulting new pathogens are commonly polylysogens, the impact of Stx prophage acquisition on the host cell’s prophage induction profile remains poorly understood.

Here, we show that the genomic carriage of a Stx prophage by the murine pathogen *C. rodentium* abrogates the induction of a co-hosted prophage during intestinal infection. Additionally, lysogenization of commensal gut *E. coli* by the infecting Stx phage results in modulation of the prophage induction profile of the newly-lysogenized bacterial cells. Moreover, we found that the Stx phage uses its Cro repressor to bind the operator of a co-hosted prophage, manipulating its regulation to inhibit phage transcription, DNA replication, and subsequent virion production. These findings uncover a novel mechanism by which an Stx phage shapes the composition of virions produced by the host cell and reveal that prophage cross-regulatory interactions play a central role in the evolution and emergence of new Stx-producing pathogens.

## Results

### Induction profile of *C. rodentium* prophages during intestinal infection

*C. rodentium* DBS100 (**CR**), a natural mouse pathogen, is a polylysogen that harbors ten prophages within its genome, which are conserved among EHEC and other enteric pathogens (*23, 24*). Of the ten prophages, five are intact, while the other five are degraded remnants (*23*). *In vitro* studies have shown that among them, ϕNP and ϕSM are the only prophages demonstrated to produce infective virions (*23, 24*).

The strain *C. rodentium* ϕ1720a-02 (**CR ϕ1720a-02**) is lysogenized with ϕ1720a-02, an Stx temperate phage originally isolated from a food-borne Stx-producing *E. coli* strain (*25*). Mouse infection with CR ϕ1720a-02 serves as an established model for STEC virulence *in vivo*, and is characterized by toxin-mediated changes in the intestinal environment, such as infiltration of phagocytes and profound inflammation (*25, 26*). Inflammatory products such as phagocytic-derived reactive oxygen species trigger prophage induction *in vitro* (*27, 28*).

We aimed to characterize the induction level of prophages in a polylysogen pathogen, referred to here as the prophage induction profile, in the context of intestinal inflammation. To this end, we infected specific pathogen free (SPF) C57BL/6 mice with either CR, CR ϕ1720a-02, or the CR ϕ1720a-02Δ*stx* (**CR ϕ1720a-02Δ*stx***) strain, in which the *stxA* and *stxB* genes in the ϕ1720a-02 prophage are deleted. We then tested whether toxin-induced inflammation affects the prophage induction profile during infection.

We collected feces throughout the course of infection and quantified the *attP* DNA site of ϕ1720a-02, ϕNP and ϕSM by quantitative PCR (qPCR). The *attP* site represents the circular, replicative form that is absent in the prophage state, thus serving as an indicator of entry into the lytic cycle (*6*). Remarkably, all three phages demonstrated similar *attP* levels in samples from both CR ϕ1720a-02 and CR ϕ1720a-02*Δstx* infected animals, indicating that prophage induction is independent of the toxin or toxin-related inflammation (**Fig. 1A-C**). Surprisingly, ϕNP induction was significantly elevated in animals infected with the CR strain compared to those infected with pathogens carrying either the ϕ1720a-02 or the ϕ1720a-02Δ*stx* prophages (**Fig. 1B**). This is in contrast to ϕSM, whose induction levels were consistent across all infection groups (**Fig. 1C**).

**Figure 1.**
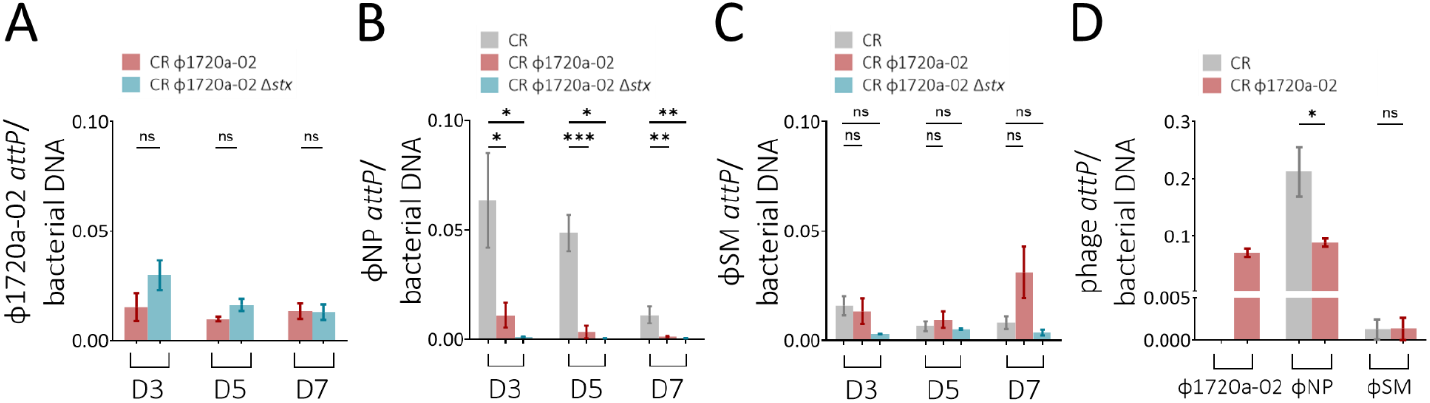
Induction profile of *C. rodentium* prophages during intestinal infection -. (A-C) Quantification of *attP* DNA levels of prophages (A) ϕ1720a-02, (B) ϕNP, and (C) ϕSM in feces of mice infected with CR, CR ϕ1720a-02, or CR ϕ1720a-02Δ*stx* at the indicated days post-infection, measured using qPCR. DNA copy numbers of the *attP* sites were determined by calibration curve and normalized to bacterial DNA. N=4-10 mice per group. (D) *attP* DNA levels of ϕ1720a-02, ϕNP, and ϕSM prophages in feces of germ-free mice (N=3) infected with CR or CR ϕ1720a-02 on day 3 post-infection, quantified by qPCR. Bars represent means ± standard error. *P<0.05. **P<0.002. ***P<0.001. ns=non-significant.

Next, we explored whether the selective induction of ϕNP observed exclusively during CR infection is driven by changes in the microbiota caused by the ϕ1720a-02 phage (*29, 30*). To investigate this, we infected germ-free mice with either CR or CR ϕ1720a-02 and quantified prophage induction in the feces of infected mice (**Fig. 1D**). As observed in conventional mice, *attP* sites were detected for all three phages, indicating that these prophages are induced in the intestine in the absence of a microbiota community. Notably, ϕNP induction was higher in germ-free mice infected with CR compared to those infected with CR ϕ1720a-02. This resembles ϕNP induction pattern in conventional mice and suggests that the gut microbiota is not responsible for the variation in ϕNP induction levels. Collectively, these findings indicate that the differences in ϕNP induction levels are associated with the presence of the ϕ1720a-02 prophage within the same bacterial host genome, rather than luminal conditions in the intestine.

### Lysogenization by ϕ1720a-02 results in modulation of the induction profile of commensal *E. coli* prophages

Next, we aimed to test whether integration of ϕ1720a-02 affects the induction of endogenous prophages following natural transmission events within bacterial strains in the gastrointestinal tract (*31, 32*). To this end, we developed an experimental assay for detecting such transmission events into resident *E. coli* strains in the mouse gut (**Fig. 2A**). We co-infected mice with CR ϕ1720a-02 Δ*stx* and a murine commensal *E. coli* strain. The toxin-deleted donor strain was used to avoid intentional generation of new STEC pathogenic strains. The ϕ1720a-02 phage carries a chloramphenicol resistance gene (*cmR*) within its lytic region (*25*). To facilitate *E. coli* recovery, we introduced a tetracycline resistance marker (*tet*) into the *lacY* gene in the lac operon. This Δ*lacY::tet* mutant exhibits no growth deficiency in culture media without lactose, nor in the gastrointestinal tract of adult mice that eat commercial standard murine diet which is lactose-free (**Fig. S1**). Selective plating was applied to verify the successful co-colonization of the murine gut by both CR ϕ1720a-02 Δ*stx* and *E. coli* (**Fig. 2B-D**). We then isolated *E. coli* Δ*lacY::tet* colonies that harbor the ϕ1720a-02 phage, representative of successful transmission events, by plating feces of infected mice on double-selective agar plates containing both antibiotics. We repeated this *in vivo* transmission assay using three different *E. coli* strains, YL151, TW482, and TW485, all of which were originally isolated from feces of healthy mice. In all co-infection sets, newly-formed lysogens were first detected two days after infection, appearing in a subset of infected animals within each group (**Fig. 2E**). From day 3 onward, we observed a decline in the carriage of newly formed lysogens, likely due to decreasing *E. coli* loads which results in low occurrence of lysogenization events dropping below the limit of detection. Carriage of ϕ1720a-02 phage by newly lysogenized *E. coli* strains was validated by PCR (data not shown). Lysogenization by ϕ1720a-02 had no impact on *E. coli* growth under standard culture conditions (**Fig. 2G-I**). One representative ϕ1720a-02 lysogen isolate from each *E. coli* strain was subjected to whole-genome sequencing and analysis, revealing that the ϕ1720a-02 phage integrated into the same chromosomal site in all *E. coli* strains, at the 5’ region of the *dusA* gene (**Fig. 2I**). The *dusA* gene encodes the tRNA-dihydrouridine synthase A and is a known site for phage integration (*33*).

**Figure 2.**
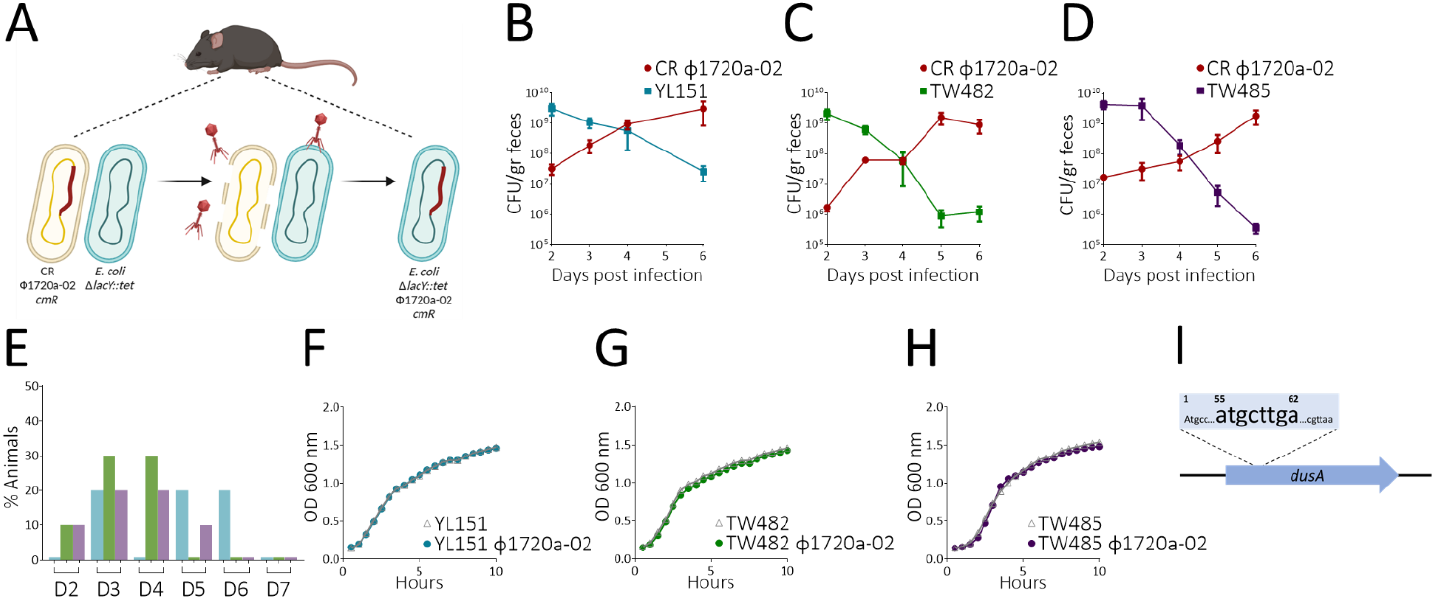
ϕ1720a-02 phage transmission in the intestine –. (A) A model illustrating the *in vivo* transmission experiment. C57BL6/J mice were co-infected with strains CR ϕ1720a-02 *cmR* and *E. coli* Δ*lacY::tet. E. coli* cells carrying ϕ1720a-02 *cmR* were isolated from feces using double-selective agar plates. (B-C) Bacterial load, measured as CFU per gram of feces, following co-infection of mice with CR ϕ1720a-02 and (B) YL151, (C) TW482, or (D) TW485 strains, determined in the course of infection by selective plating. (E) The percentage of mice shedding *E. coli* Δ*lacY::tet* ϕ1720a-02 *cmR* in each infection group throughout the course of infection. (F-H) Growth curves of strains (F) YL151, (G) TW482, and (H) TW485, along with their respective ϕ1720a-02 lysogens, in LB media. (I) Schematic representation of the ϕ1720a-02 integration site, within the 5’ region of *C. rodentium dusA* gene. The numbers indicate the base positions within the coding sequence, the enlarged sequence highlights the specific integration region.

To investigate how ϕ1720a-02 lysogenization affects the host cell prophage induction profile, we used long-read sequencing and coverage analysis of uninduced and induced bacterial cultures, as done in previous studies (*34, 35*). Briefly, following induction, both phage and bacterial DNA were purified and subjected to Oxford Nanopore sequencing. Putative prophage regions in the bacterial genome showing increased coverage compared to mean chromosomal coverage are indicative of elements undergoing active phage DNA replication (**Fig. 3**).

**Figure 3.**
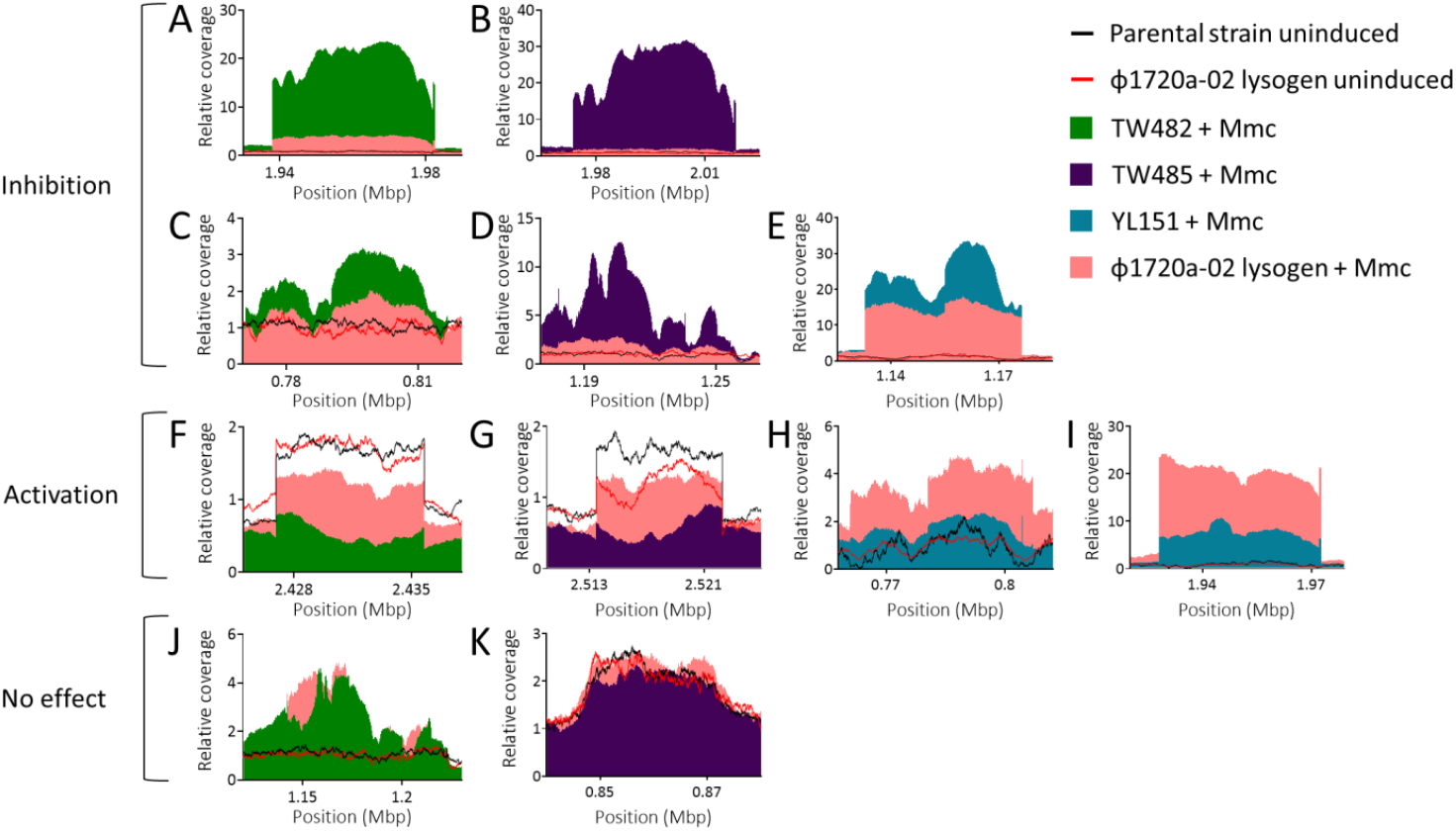
Lysogenization by ϕ1720a-02 results in modulation of the induction profile of commensal *E. coli* prophages. Relative coverage of phage genomic DNA and proximal chromosome regions in commensal *E. coli* strains and their respective ϕ1720a-02 lysogen strain. Phage and bacterial DNA were purified from uninduced and mitomycin C (MmC)-treated cultures and subjected to Oxford Nanopore sequencing. Coverage of untreated samples is represented by lines, while coverage of treated samples is displayed as a shaded area graph. Coverage higher than the coverage in chromosome regions is indicative of active phage DNA replication.

The parental strains YL151, TW482, and TW485 carry 7, 5, and 6 prophages, respectively, including intact and incomplete prophages. Of these, we identified 3-4 prophages in each strain exhibiting varying degrees of active DNA replication (**Fig. 3**). Notably, lysogenization with ϕ1720a-02 resulted in significant alterations in the induction levels of several prophages. Specifically, the most highly replicative prophages in the parental strains exhibited either substantial (**Fig. 3A-B**) or moderate (**Fig. 3C-E**) inhibition of induction in the ϕ1720a-02 lysogen. Conversely, some prophages that were only mildly induced in the parental strains showed increased induction levels in the ϕ1720a-02 lysogen (**Fig. 3F-I**), with one showing strong induction in the lysogen (**Fig. 3I**). Interestingly, certain prophages exhibited no change in induction levels between the parental strains and the ϕ1720a-02 lysogen, regardless of whether induction was triggered by treatment (**Fig. 3J**) or occurred spontaneously (**Fig. 3K**). Three prophages showed mild induction in the absence of a trigger, in both the parental and the ϕ1720a-02 lysogen (**Fig. 3F-G, K**)

Taken together, these data suggest that the ϕ1720a-02 phage produces infective virions that transmit to and lysogenize new commensal *E. coli* hosts in the intestine, thereby highlighting its potential to create new pathogens. Importantly, ϕ1720a-02 integration into the host chromosome significantly alters its prophage induction profile. These effects may occur either directly, through ϕ1720a-02 interfering with the induction of co-hosted prophages, or indirectly, by interfering with an element that itself modulates the induction of other co-hosted prophages.

### ϕNP induction is inhibited when co-hosted with ϕ1720a-02

Having established that ϕ1720a-02 interferes with activity of other co-hosted prophages, we next aimed to identify the specific stage at which it disrupts the induction process. To this end, we investigated the interaction between ϕ1720a-02 and ϕNP in CR, as it provided a simplified host model in which only a single prophage was induced per strain. To test the ability of *C. rodentium* native prophages to form infective virions, we used mitomycin C to trigger prophage induction in *C. rodentium* strains, filtered the virions released into the supernatant, and infected a lawn of susceptible *E. coli*. Notably, ϕ1720a-02 phage does not form visible plaques (*6*). Thus, the observed plaques inevitably result from infective virions of the other encoded prophages (**Fig. 4A**). Virions produced by CR ϕ1720a-02 generated fewer visible plaques compared to those produced by CR, indicating that ϕ1720a-02 carriage reduces the ability of its host cell to produce plaque-forming virions encoded by co-hosted prophages. This was independent of the Shiga toxin, as virions produced by CR ϕ1720a-02Δ*stx*, demonstrated a similar hindrance in the production of plaque-forming virions.

**Figure 4.**
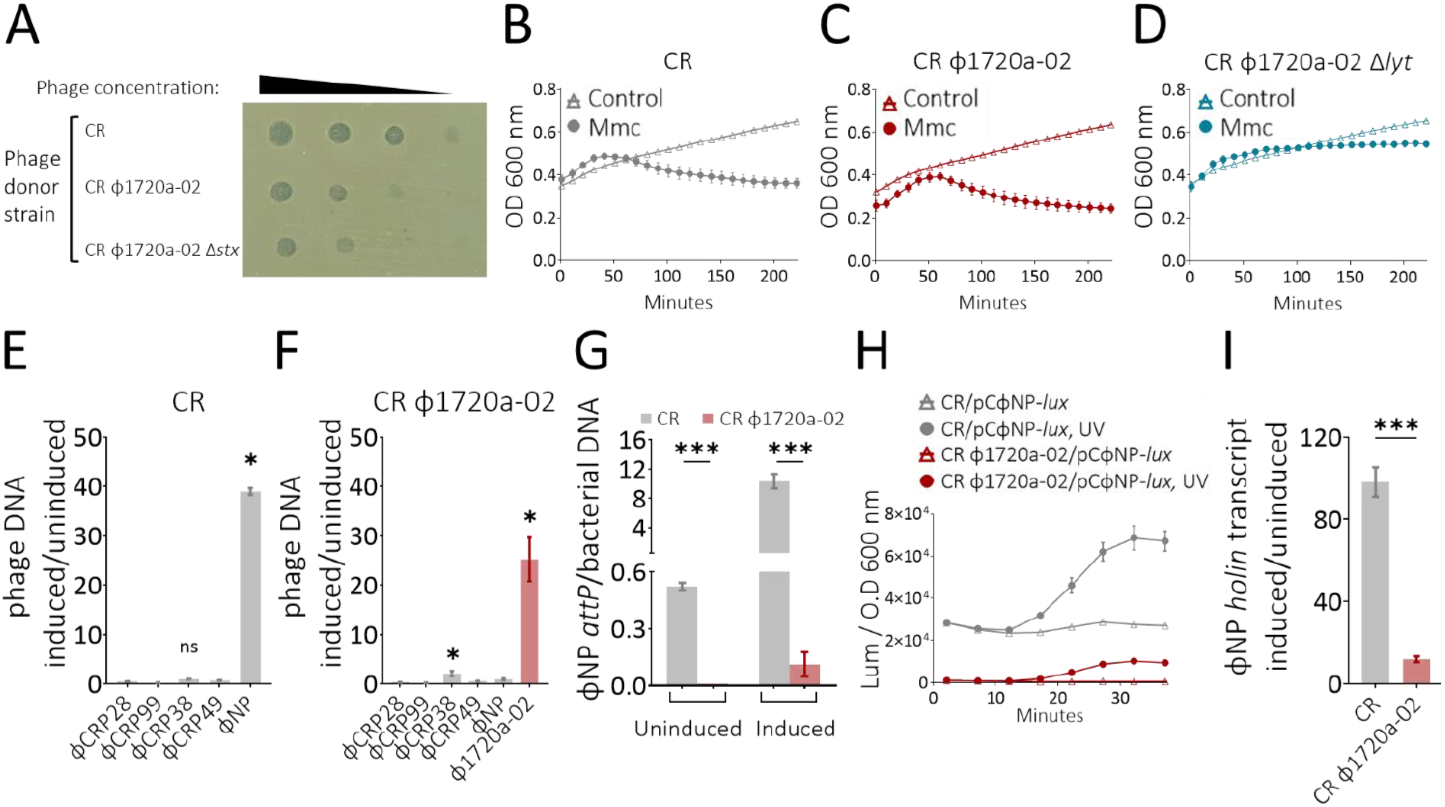
ϕNP induction is inhibited when co-hosted with ϕ1720a-02. (A) Plaques formed on an *E. coli DH5α* lawn by phages produced by the indicated donor strains treated by mitomycin C (MmC). Phages were serially diluted 10-fold. (B-D) Growth curves of (B) CR (C) CR 1720a-02 (D) CR 1720a-02 Δ*lyt* strains in LB broth, treated with MmC, or untreated. (E-F) DNA levels of prophages encoded by (E) CR or (F) CR ϕ1720a-02 measured by qPCR. DNA levels are presented as relative copies in MmC-treated vs. untreated cultures, normalized to bacterial DNA. (G) DNA levels of ϕNP *attP* site of uninduced or induced cultures of the indicated strains, measured by qPCR, DNA levels were quantified by calibration curve and normalized to bacterial DNA. (H) Transcription of the early lytic operon of ϕNP in untreated and UV-treated cultures. The strains carry pCϕNP-lux, encoding the control region of the operon positioned upstream of a lux reporter. Data is presented as luminescence normalized to OD600nm. (I) Transcript level of holϕNP RNA of the indicated strains presented as relative copies in UV-treated vs. untreated cultures, normalized to *rpoA* transcript. Strains used are CR, CR ϕ1720a-02, CR ϕ1720a-02Δ*stx* deleted in toxin genes, CR ϕ1720a-02Δ*lyt*, deleted in the lytic genes of ϕ1720a-02 prophage. Bars represent means ± standard error. *P<0.05. **P<0.002. ***P<0.001. ns=non-significant.

Next, we assessed the ability of the encoded prophages to trigger lysis in each host cell. Uninduced bacterial strains CR and CR ϕ1720a-02 demonstrated similar growth under standard culture conditions (**Fig.4B-D**). Treatment with mitomycin C led to a rapid decline in optical density in both cultures, a characteristic pattern indicative of bacterial cells undergoing prophage-mediated lysis. To distinguish between ϕ1720a-02-induced lysis and lysis by the other *C. rodentium* prophages, we generated a mutant strain deleted in the ϕ1720a-02 lytic genes (**CR ϕ1720a-02Δ*lyt***). Markedly, following treatment with mitomycin C, the growth of CR ϕ1720a-02Δlyt stagnated, but no reduction in optical density was observed, suggesting that lysis, as seen in CR, did not occur. This suggests that the other encoded prophages are unable to trigger cell lysis in a cell that is lysogenized with ϕ1720a-02.

Cell lysis and virion production in a bacterial cell are preceded by transcription of lytic phage genes and phage DNA replication which are the initial events following entry into the lytic life cycle. We next investigated phage DNA replication following mitomycin C induction by qPCR. The CR strain showed significant DNA replication of ϕNP, but not of other prophages, compared to untreated samples (**Fig. 4E**). However, the CR ϕ1720a-02 strain exhibited significant replication of the ϕ1720a-02 phage, and mild replication of ϕSM, but almost no replication of ϕNP (**Fig. 4F**). This selective induction pattern was similar when prophage induction was triggered by UV irradiation, a known trigger of the SOS response (**Fig. S2**). We then quantified ϕNP DNA excision by qPCR of its *attP* site and found that upon induction, excision was significantly reduced in the CR ϕ1720a-02 strain compared to CR (**Fig. 4G**). Additionally, basal level of ϕNP DNA excision without any induction trigger, which naturally occurs at low frequency, was abrogated in the CR ϕ1720a-02 strain (**Fig. 4G**). This suggests that ϕ1720a-02 disrupts ϕNP DNA excision and replication both in the lysogenic state and upon activating the lytic cycle. These findings corroborate our observation of lower ϕNP phage DNA excision and replication in mice infected with CR ϕ1720a-02 compared to those infected with CR (**Fig. 1B**).

To study transcription of lytic ϕNP genes, we constructed a reporter plasmid that harbors the ϕNP control region, spanning between the CI repressor and the Cro repressor of ϕNP, upstream to the *lux* reporter operon (pCϕNP-*lux*) (*36*). This reporter plasmid was then transformed into CR, and luminescence was monitored as an indicator of the early lytic ϕNP gene expression. Basal expression level of ϕNP lytic genes was significantly lower in the CR ϕ1720a-02 strain in comparison with expression level in CR (**Fig. 4H**). Furthermore, upon induction, the increase in early ϕNP gene expression observed in CR was significantly higher than that in CR ϕ1720a-02. Additionally, we used quantitative reverse-transcription PCR to measure transcript levels of the *hol* gene of ϕNP (*hol*_*ϕNP*_), which encodes Holin - a key protein involved in bacterial membrane disruption and facilitates phage-induced lysis. The *hol*_*ϕNP*_ transcript levels were significantly higher in induced CR compared to induced CR ϕ1720a-02 (**Fig. 4I**). We conclude that ϕ1720a-02 inhibits the transcription of ϕNP lytic genes, blocking the initial stages of the switch from the lysogenic to the lytic cycle.

These findings demonstrate that the Stx phage, ϕ1720a-02, inhibits early lytic gene expression, DNA replication, DNA excision, phage-mediated lysis and virion production of co-hosted ϕNP phage. The observation that ϕ1720a-02 modifies the baseline functions of ϕNP prophage demonstrates that ϕ1720a-02 cross-regulates prophage activities within lysogenic host cells under non-inducing conditions as well.

### Cro_Φ1720a-02_ represses the induction of co-hosted ϕNP

Next, we sought to uncover the mechanism by which the ϕ1720a-02 phage inhibits the induction of ϕNP. The inhibition of ϕNP lytic operon gene expression in strains co-hosted with ϕ1720a-02 suggests that ϕ1720a-02 interferes with ϕNP induction at the transcriptional level, possibly through a transcriptional regulator acting *in trans* (**Fig. 4H-I**). The first factor encoded in the lytic operon of lambdoid phages, including Stx phages, is the Cro repressor. Cro drives the transition into the lytic cycle by repressing the activity and expression of CI repressor, responsible for maintaining lysogeny. A previous report showed that the Cro factor of an Stx phage in EHEC modulates the expression of various genes outside of the phage genome (*37*). This led us to test whether the Cro of ϕ1720a-02 affects gene expression of co-hosted prophages. To this end, we constructed **CR ϕ1720a-02Δ*cro***, a variant strain deleted in its *cro* repressor gene. The Cro protein is essential for repressing CI, the lysogeny-maintenance gene, and activating lytic genes. Deleting the *cro* gene in a prophage disrupts its ability to transition from lysogeny to the lytic cycle (*38*). As expected, this deletion abolished ϕ1720a-02 replication (**Fig. 5A**). In parallel, it also restored the replication ability of the ϕNP prophage in CR ϕ1720a-02 (**Fig. 5B**), as well as its DNA excision and circularization (**Fig. 5C**). Complementation of c*ro*_Φ*1720a-02*_ via an expression plasmid (pBAD-*cro*_Φ*1720a-02*_) resulted in the abrogation of ϕNP replication upon induction, an effect not observed with the control plasmid (pBAD, **Fig. 5D**). Similarly, *cro*_Φ*1720a-02*_ complementation, but not the control plasmid, protected CR ϕ1720a-02Δ*cro* from phage-mediated lysis (**Fig. 5E-F**).

**Figure 5.**
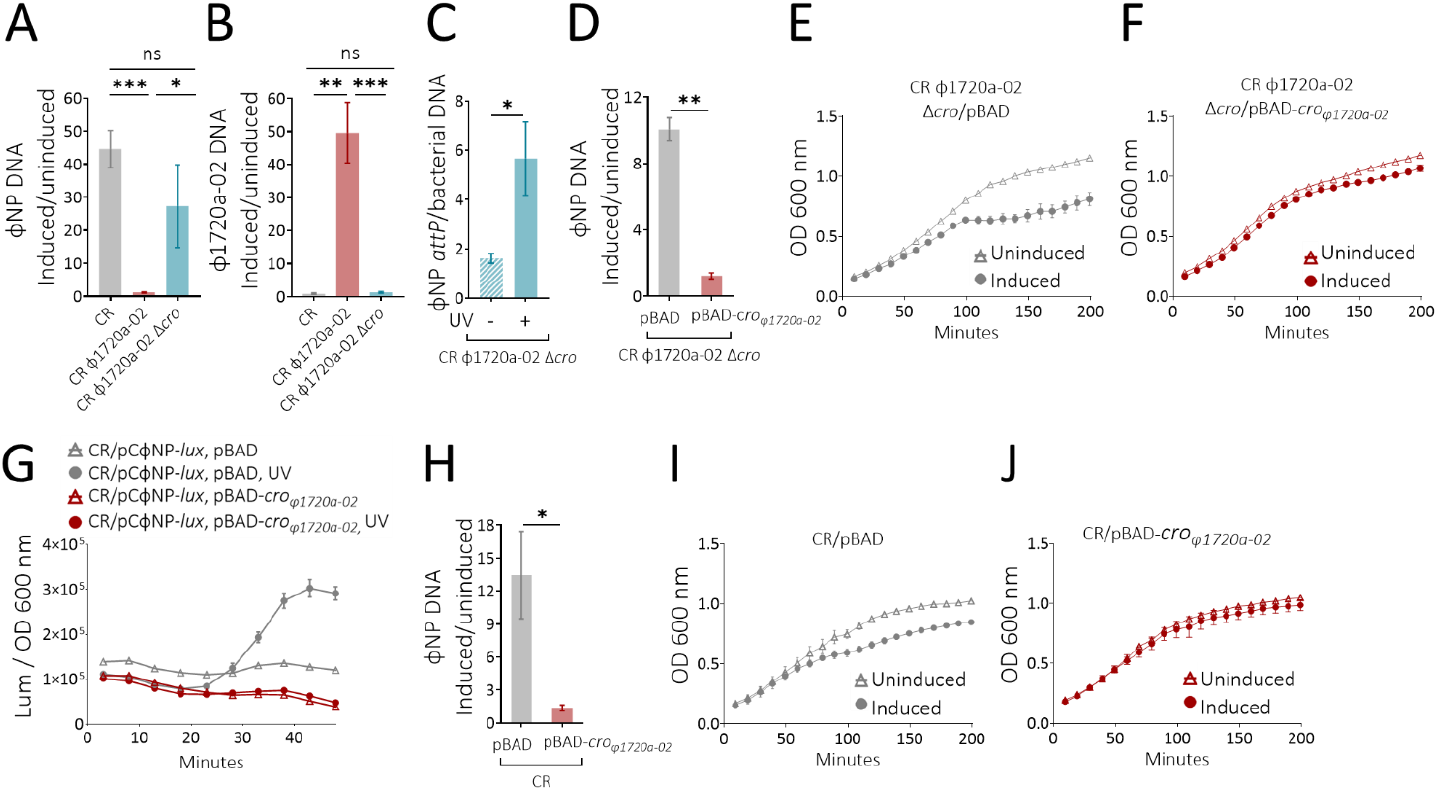
Cro1720a-02 represses the induction of co-hosted ϕNP. – (A-B) DNA levels of (A) ϕNP and (B) ϕ1720a-02 measured by qPCR in CR, CR ϕ1720a-02, or CR ϕ1720a-02Δ*cro* deleted in the *cro* repressor gene. DNA levels are presented as relative copies in UV-treated cultures vs. untreated cultures, normalized to bacterial DNA. (C) Copy numbers of ϕNP *attP* DNA in uninduced or UV-induced cultures of CR ϕ1720a-02Δ*cro*, measured by qPCR, normalized to bacterial DNA copy numbers. (D) DNA levels of ϕNP measured by qPCR in CR ϕ1720a-02Δ*cro* harboring a plasmid encoding the Cro_1720a-02_ repressor, or the control plasmid pBAD. DNA levels are presented as relative copies in UV-treated cultures vs. untreated cultures. DNA levels were quantified by calibration curve and normalized to bacterial DNA. (E-F) Growth curves of untreated or UV-treated CR ϕ1720a-02 Δ*cro*, complemented with (E) the control plasmid (F) the Cro_1720a-02_ repressor. (G) Cro1720a-02 effect on transcription of the early lytic operon of ϕNP in untreated and UV-treated cultures. All presented strains harbor a plasmid encoding the Cro1720a-02 repressor, or a control plasmid, in addition to the pCϕNP-lux reporter plasmid, which contains the control region of the operon positioned upstream of a lux reporter. Data is presented as luminescence normalized to OD600nm in media supplemented with arabinose. (H) Cro1720a-02 effect on DNA replication of ϕNP measured by qPCR in CR strains harboring a plasmid encoding the Cro1720a-02 repressor, or a control plasmid, in media supplemented with arabinose. (I-J) Growth curves of untreated or UV-treated CR, harboring (I) the control plasmid pBAD or (J) a plasmid encoding Cro1720a-02 repressor. Bars represent means ± standard error. *T-test P<0.05. **T-test P<0.002. ***T-test P<0.001. ns=non-significant.

Next, we studied whether Cro_Φ1720a-02_ is sufficient to hinder ϕNP induction in CR, within a host that does not carry the ϕ1720a-02 genome. For this, we transformed the CR*/*pCϕNP*-lux* reporter strain with either pBAD- *cro*_*Φ1720a-02*_ or the control plasmid pBAD, and monitored ϕNP lytic gene expression. As expected, upon induction, the reporter strain with the pBAD control plasmid showed a significant increase in ϕNP lytic operon expression (**Fig. 5G**). Markedly, the strain harboring pBAD-*cro*_*Φ1720a-02*_ exhibited no such increase in expression levels, indication that induction was inhibited by Cro_Φ1720a-02_ (**Fig. 5G**). Moreover, ϕNP DNA replication was significantly inhibited in CR*/*pBAD-*cro*_*Φ1720a-02*_ compared to the replication observed in CR*/*pBAD (**Fig. 5H**). Additionally, Cro_1720a-02_ expression in CR abolished phage-mediated bacterial lysis following prophage induction (**Fig. 5I-J**). Since ϕNP is the only actively replicating phage in CR (**Fig. 4E**), we attribute the lysis to ϕNP induction and conclude that the expression of Cro_Φ1720a-02_ protected CR from ϕNP-mediated lysis.

Our findings demonstrate that the Cro repressor of ϕ1720a-02, not only promotes its own lytic cycle, but also unexpectedly inhibits the lytic cycle of the co-hosted ϕNP. This demonstrates how a phage-encoded factor can exert opposing regulatory effects on co-hosted prophages, inhibiting their induction and reducing competition for host cell resources.

### Structural modeling predicts dual function for Cro_Φ1720a-02_

Cro repressors are small DNA-binding proteins typically found in temperate bacteriophages, where they play a crucial role in the lysis-lysogeny decision switch. Following induction and CI degradation, Cro repressors are expressed and promote the lytic cycle by binding to operator sites, preventing further CI transcription. This alleviates repression of the lytic operon and initiates the lytic cycle. Functioning as dimeric structures, Cro repressors feature a distinctive compact fold that includes a helix-turn-helix (HTH) motif. This HTH motif enables Cro repressors to recognize and bind to specific operator sequences within the DNA (*39*). Despite sharing conserved secondary structure elements and a similar overall fold, Cro proteins show considerable diversity in how they interact with their operator sequences (*40, 41*). This variability exists both in the amino acids that contact the DNA and in the nucleotide sequences of the operator regions themselves, thus enabling phage-specific regulation of the induction process (*41*).

We utilized AlphaFold3 to predict the structure of Cro_Φ1720a-02_ (*42*). The predicted structure showed high similarity to the crystal structure of the Cro protein from the P22 phage of *Salmonella enterica* (**Fig. 6A**). The predicted structure of Cro_Φ1720a-02_ comprises of five helices within the 67-residue N-terminal region, corresponding to the DNA-binding domain of P22 Cro, and an unstructured C-terminal tail typical to this family of proteins.

**Figure 6.**
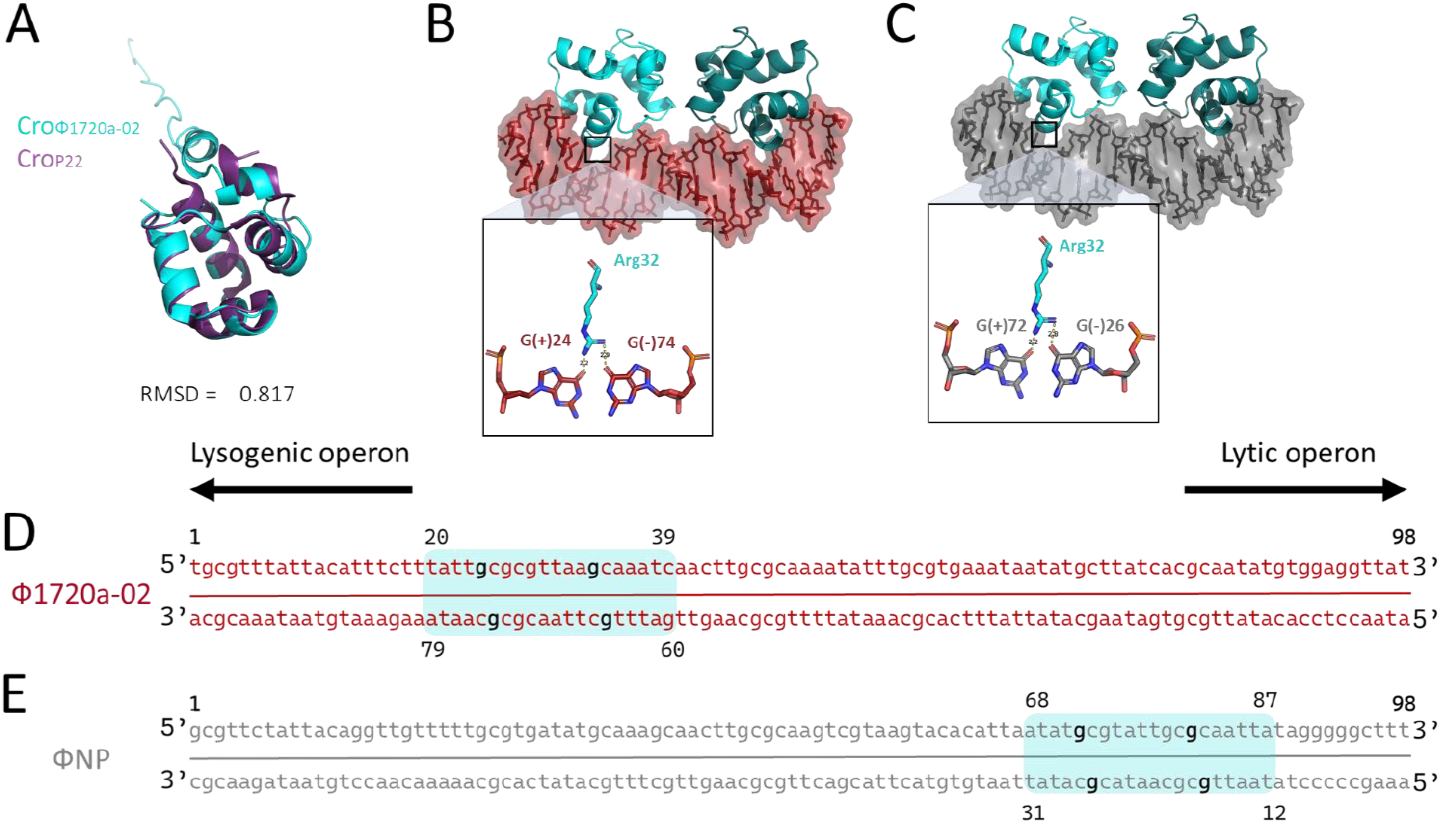
Structural modeling predicts dual function for Cro1720a-02 –. (A) Structural alignment of the Cro1720a-02 monomer AlphaFold3 prediction with the 2R1J crystallography structure of Crop22, visualized using PyMOL. (B) Structural model of the Cro_1720a-02_ dimer bound to its predicted DNA-binding site within the ϕ1720a-02 control region. The inset (square) illustrates the specific hydrogen bonds formed between Cro Arginine 32 and guanine bases at positions (+)24 and (−)74 of the ϕ1720a-02 binding site. (C) Structural model of the Cro1720a-02 dimer bound to its predicted DNA-binding site within the ϕNP control region. The inset (square) illustrates the specific hydrogen bonds formed between Cro arginine 32 and guanine bases at positions (+)72 and (−)26 of the ϕNP binding site. (D-E) Illustration of the (D) 98 bp ϕ1720a-02 control region sequence and (E) 98 bp ϕNP control region sequence. The 20 bp Cro predicted binding site is highlighted in blue, with numbers indicating its position within each control region. The guanine bases involved in specific hydrogen bonding with Cro arginine 32 are shown in bold. The direction of the downstream lytic and lysogenic operons is indicated by arrows.

We then used AlphaFold3 to model the dimeric interaction of the Cro_Φ1720a-02_ DNA-binding region with the entire control region of ϕ1720a-02. The modeling analysis predicted that the Cro_Φ1720a-02_ dimer binds to a potential operator sequence - a 20 base pair region located 20-39 bases upstream of ϕ1720a-02’s lysogenic operon. The inter-chain predicted Template Modeling (ipTM) score of 0.92 indicates strong confidence in the predicted binding interface (**Fig. 6B, 6D)** (*43*). Using PDBePISA (*44*) we identified a specific interaction where Arg32 of Cro_Φ1720a-02_ forms hydrogen bonds with two guanine residues, one on each strand of the DNA. These guanine residues are located within the semi-palindromic sequence, and we speculate that they, along with their interactions with arginine, play a central role in determining the interface specificity.

Next, we aimed to determine whether there is any predicted interaction between the Cro_Φ1720a-02_ and the control region of ϕNP. Remarkably, AlphaFold3 predicted, with high confidence (ipTM=0.92), that the Cro_Φ1720a-02_ DNA-binding domain also interacts with the control region of ϕNP. In this case, this putative operator of 20 bases resides 12 bases upstream of the first gene in the ϕNP lytic operon, explaining its inhibitory effect on ϕNP lytic gene transcription **(Fig. 6C, E**). This predicted Cro_Φ1720a-02_-binding sequence contains the same two guanine residues, spaced by eight bases, as in the ϕ1720a-02 control region. These guanines are predicted to form hydrogen bonds with Cro_Φ1720a-02_ Arg32, identical to the bonds observed in the structure of Cro_Φ1720a-02_ bound to the ϕ1720a-02 sequence.

This data suggests that binding of the Cro_Φ1720a-02_ repressor to the control regions of the phages exerts different outcomes – repressing lysogeny for ϕ1720a-02 and repressing induction of the lytic life cycle in ϕNP. Despite high sequence variability among Cro repressors (*45*), we identified 16 Stx phages, including those isolated from infected patients and livestock carriers, that encode a Cro protein with identical or highly similar sequences to Cro_Φ1720a-02_, all of them contain a conserved arginine in position 32 (**Fig. S3**). The Cro repressor encoded by these Stx phages may also play a role in controlling the activity of the other prophages present within these STEC strains.

Overall, our structural modeling of Cro_Φ1720a-02_ revealed its ability to form dimeric interactions with the control regions of both ϕ1720a-02 and ϕNP. The findings demonstrate Cro’s dual functionality: promoting the lytic cycle of its own encoding phage while repressing the expression of ϕNP lytic genes.

## Methods

### Mouse Experiments

All animal studies were approved by the Authority for Biological and Biomedical Models of the Hebrew University and comply with the International Animal Care and Use Committee (IACUC) guidelines. Animals were housed in cages in a temperature-controlled room at 22°C and 12-hour light/dark cycle with ad libitum access to food and water and inspected regularly by SPF personnel.

### C. rodentium infection

5-8-week-old specific pathogen free (SPF) C57BL/6J mice (Envigo #043) were randomly assigned into individually ventilated cages and allowed to acclimate in our ABSL2 vivarium for at least 5 days undisturbed. For infections with C. rodentium strains, mice were pre-treated with an intragastric dose of 2.5 mg vancomycin in 100 ul DDW. 24 hours later, mice were inoculated with 5×10^9^ CFU per mouse of a single *C. rodentium* strain suspended in 100 μl PBS via oral gavage. Bacterial load in feces and dosage of bacterial inoculum were determined by serial dilution and plating on selective MacConkey agar plates.

### E. coli and C. rodentium co-infection

5-8-week-old SPF C57BL/6J mice were pre-treated with an intragastric dose of 20mg streptomycin in 100ul DDW. Three hours later mice were inoculated with 5×10^9^ CFU per mouse of a single *E. coli* strain suspended in 100μl PBS via oral gavage. 24 hours following *E. coli* infection, mice were inoculated with 5×10^9^ CFU per mouse of a single C. rodentium strain suspended in 100μl PBS via oral gavage. Bacterial load in feces and dosage of bacterial inoculum were determined by serial dilution and plating on selective MacConkey agar plates.

### Germ free mice infections

Germ-free C57BL/6J mice were bred and housed at the germ-free facility of the Authority for Biological and Biomedical Models at The Hebrew University. At six weeks of age, the mice were aseptically removed from the isolator, transferred to a biosafety cabinet, and placed into autoclaved, individually ventilated cages. Infections were performed within the biosafety cabinet, where mice were inoculated with 5 × 10^7^ CFU of a single *C. rodentium* strain suspended in 200 μl of PBS per mouse via oral gavage. Bacterial load in feces and dosage of bacterial inoculum were determined by serial dilution and plating on selective MacConkey agar plates.

### Plaque-forming assay

Overnight *E. coli* DH5α bacterial cultures were mixed with MB 0.5% agar (LB medium supplemented with 0.1 mM MnCl_2_, 5 mM MgCl_2_, 5 mM CaCl_2_, 5 μM NaOH, and 0.5% agar), poured onto MB 1.6% agar plates, and left to solidify. Prophage induction was performed by growing the indicated strains to an OD_600_ of 0.4, followed by treatment with 0.5 μg/mL mitomycin C for 2 hours at 37°C with shaking at 220 rpm. After 2 hours, cells were centrifuged at 14,000 rpm for 1 min, and the supernatant was filtered through a 0.45 μm filter. Phage filtrates were serially diluted tenfold and 5 μL from each dilution were spotted onto the MB agar plates embedded with target bacteria. The plates were incubated overnight at 37°C.

### Bacterial strains

Bacterial strains used in this study are listed in Table S1. Bacterial strains were routinely grown in LB broth or on LB-agar plates unless otherwise indicated. Antibiotics were used at the following concentrations: ampicillin (Amp), 0.1 mg/ml; chloramphenicol (Cm), 0.025mg/ml; kanamycin (Kan), 0.05 mg/ml; tetracycline (Tet), 0.01 mg/ml; streptomycin (strep), 0.1mg/ml. Plasmids were propagated in *E. coli* strain DH5α λpir or MFD λpir. The MFD λpir strain was grown in LB medium supplemented with 0.3 mM diaminopimelic acid.

### Plasmid design and construction

Plasmids used in this study are listed in Table S2. Plasmids were constructed using the Gibson Assembly method. DNA fragments were designed with overlapping sequences at their ends, typically ranging from 20 to 40 bp. The Gibson Assembly reaction was carried out in a 20 µL mixture containing 100 ng of the vector DNA fragment, with non-vector fragments calculated based on NEB instructions and added in the correct amounts, along with homemade Gibson Assembly. The reaction was incubated at 50°C for 1 hour.

pBAD*-croϕ1720a-*02 (pRU517) was constructed as follows: PCR fragments from pRU514 were amplified with primers 643/644, followed by a second PCR with primers 835/836 to add an RBS and x6HIS tag. The *cro* gene was amplified from DBS770 with primers 837/838. The products were assembled using Gibson Assembly. p*C*_*ϕNP*_*-lux* (pRU575) was constructed as follows: The C5-5 promoter in pRU565 was replaced with the ϕNP lytic control region by PCR amplification of pRU565 with primers 885/886, followed by circularization using Gibson Assembly.

pACYC184-*C*_*ϕNP*_*-lux* (pRU654) was constructed as follows: The pACYC184 backbone was amplified with primers 940/941. The lux operon with the ϕNP lytic operator was amplified with pRU575 as template with primers 894/939. The fragments were assembled using Gibson Assembly.

### Construction of bacterial mutants

The Δ*lyt::kan*, Δ*cro::kan*, Δ*stxAB::kan* (labeled Δ*lyt*, Δ*cro*, Δ*stx*, respectively), and Δ*lacY::tet* mutants were constructed using the lambda red recombination system and selective cassettes (*46*). Briefly, *C. rodentium* or *E. coli* strains carrying the pKD46-strep plasmid were grown at 30°C until they reached OD600nm 0.2 and then recombinase expression was induced by adding 10 mM L-arabinose. Bacteria were harvested at OD600nm 0.6 and then electroporated with a PCR fragment that contains a resistance cassette with flanking regions that are homologous to the target gene. Plasmids and primers used for cloning are listed in Tables S2 and S3, respectively.

To generate Δ*lacY::tet E. coli* mutants, a *tet* cassette was PCR amplified using 323 and 324 primers containing ∼100 bp homologous sequences flanking the upstream and downstream regions of *lacY* ORF. The target fragment containing the tet cassette was then introduced into the *E. coli* strain using the λ Red recombination system, as described above.

To generate Δ*lyt::kan* mutants, a *kan* cassette was PCR amplified using primers 403 and 404 containing ∼60 bp homologous sequences flanking the upstream and downstream regions of a ∼6700 bp region in the 1720α-02 genome, which encodes the lytic genes. The target fragment containing the *kan* cassette was then introduced into the *C. rodentium* strain using the λ Red recombination system, as described above.

To generate Δ*cro::kan* and Δ*stxAB::kan* mutants, fragments flanking each gene and the *kan* cassette were cloned into plasmids pRU533 (pEP185.2-*cro::kan*) or pTW475 (pUC19-*stxAB::kan*), which were used as the template to generate the target fragments for recombination.

pTW475 was constructed as follows: 1kb homologous regions upstream and downstream of *stxAB* were PCR-amplified from DBS770 with primers 738/739 and 740/741, respectively. The pUC19 backbone was amplified with primers 485/486, and the *kan* cassette with primers 254/255. All fragments were assembled using Gibson Assembly.

plasmids pRU533 was constructed as follows: 1kb homologous regions upstream and downstream of *cro* were PCR-amplified from DBS770 with primers 869/870 and 871/872. The pEP185.2 backbone and *kan* cassette were amplified with primers 578/579 and 513/514, respectively. All fragments were assembled using Gibson Assembly.

Mutant genotypes were routinely confirmed by PCR. All knockout mutations were designed with the antibiotic resistance cassette inserted in the inverse orientation relative to the bacterial open reading frames (ORFs) to minimize polar effects.

### Bacterial growth assay

The indicated strains were grown at 37°C until reaching an OD600nm ≈ 0.6. When indicated, cultures were treated with UV light for 10 seconds using a biological hood lamp or with 0.5 μg/mL mitomycin C for 30 minutes. After mitomycin C treatment, cells were washed twice with PBS to remove residual mitomycin C and re-cultured in fresh LB medium. When indicated, 5 mM L-arabinose was added when the cultures reached an OD600nm ≈ 0.2. Cells were then transferred to a 96-well plate and incubated in a plate reader at 37°C with continuous shaking (double orbital motion at 307 cpm, 5 mm orbit). OD600nm measurements were recorded every 10 minutes.

### Measurements of bioluminescence

Bioluminescence was measured in strains carrying the plasmids p*C*_*ϕNP*_*-lux* or pACYC184-*pC*_*ϕNP*_*-lux*. The indicated strains were grown at 37°C until reaching an OD600nm ≈ 0.6. The indicated cultures were then treated with UV light for 10 seconds using a biological hood lamp. When indicated, 5 mM L-arabinose was added at OD600nm = 0.2. Cells were then transferred to a 96-well plate and incubated in a plate reader at 37°C with continuous shaking (double orbital motion at 307 cpm, 5 mm orbit). OD600nm measurements and bioluminescence measurements were recorded every 5 minutes.

### RNA Purification and cDNA Synthesis

RNA was extracted from bacterial cultures using the NucleoSpin RNA kit (MN, MAN-740955.50) following the manufacturer’s protocol. Complementary DNA (cDNA) was synthesized from the purified RNA using the High-Capacity cDNA Reverse Transcription Kit (ABI, AB-4374966). qPCR was performed as detailed below.

### Quantitative PCR

The indicated strains were grown and treated as described in the text and according to the “Bacterial Growth Assay” method. Following treatment, cells were re-incubated for 2 hours at 37°C with shaking at 220 rpm, then heat-treated at 95°C for 5 minutes. Debris was removed by centrifugation at 13,000 rpm for 1 min, and 1 μL of the lysate was used as the DNA template. qPCR was performed using the Luna Universal qPCR Master Mix (NEB, M3003E) following the manufacturer’s protocol. Reaction was carried out on a QuantStudio machine and fluorescence data were collected at the end of each cycle, and melting curve analysis was performed to confirm specificity. Quantification of qPCR results was performed using two methods. For relative quantification, the ΔΔCT method was used to compare treated and untreated samples, with normalization to the *eae* gene as a bacterial genomic DNA control. For absolute quantification of *attP*, a calibration curve was generated using serial dilutions of known DNA concentrations. All data were normalized to *eae* to account for variations in bacterial genomic DNA levels.

### Whole genome sequencing and coverage analysis

Genomic sequencing of the *E. coli* strains was performed using Oxford Nanopore Technology with custom analysis and annotation. An amplification-free long-read sequencing library was constructed using the v14 library prep chemistry, ensuring minimal DNA fragmentation. The library was sequenced with R10.4.1 flow cells using a primer-free protocol, generating raw data in FASTQ format. Raw reads were filtered and improved in quality using Filtlong (v0.2.1). The genome was initially assembled with Miniasm (v0.3) to create a rough sketch, followed by a high-quality assembly with Flye (v2.9.1) (*47, 48*). The Flye assembly was polished using Medaka (v1.8.0). Reads were subsequently mapped to the genome using Minimap2 by Geneious Prime 2025.0.3 with default settings, with secondary alignments excluded to enhance mapping clarity. To normalize sequencing coverage across experiments, the average coverage was calculated along the bacterial chromosome, excluding prophage regions. Each base coverage, including prophages, was then divided by the average genome coverage determined in the previous step for each experiment. Final normalized coverage graphs were generated and visualized using Prism.

### Data visualization and statistical analysis

Data displaying bacterial numbers (CFU/g) or bar graphs represent the mean and the standard error of the mean. Significance of differences was determined by Mann-Whitney non-parametric test for animal experiments and unpaired T-test for *in vitro* experiments. A P-value of less than 0.05 was considered significant. Grubb’s test was performed to detect and eliminate outlier data points.

## Discussion

Pathogen genomes are notably enriched with prophages that contribute significantly to pathogen fitness (*49*). For example, EHEC carries multiple prophages that encode virulence factors, including type III secretion system effectors, which are essential for successful colonization and infection (*19, 50*). An additional advantage to bacterial fitness comes from prophage-encoded superinfection exclusion (SIE) factors, each protecting against specific viral invaders. Thus, preserving diverse prophages within a genome equips the host cell with broad immunity (*51*). Consequently, selective pressure favors the retention of multiple prophages within the pathogen’s genome.

Prophages encoded within a polylysogen host compete with their counterparts for successful replication and transmission. This competition becomes critical when the bacterial SOS response triggers prophage induction, committing the cell to lysis and disabling further replication of the prophages as part of the bacterial chromosome. During this race against cell death, each prophage must replicate independently of the chromosome and assemble infectious virions to ensure its survival.

Only a handful of mechanisms are known by which a specific prophage can outcompete other co-hosted prophages during induction. One such mechanism is exhibiting a shorter lysis time, which is associated with higher virion productivity and provides a competitive advantage among co-hosted prophages (*52*). In the soil bacteria *Bacillus thuringiensis*, a prophage-encoded protein binds the bacterial SOS response repressor LexA, thus preventing the induction of a competing prophage, and allowing its own prophage to replicate first (*53*). Some prophages gain an advantage by evolving sensitivity to distinct environmental signals, allowing them to induce when other prophages remain dormant, thereby avoiding competition (*54*). Here, we describe a newly identified competitive strategy in which an Stx phage uses its Cro repressor to directly suppress the induction of a co-hosted prophage while driving its own lytic cycle.

Sharing evolutionary origins, the CI and Cro repressors evolved to play opposing roles in the same phage regulatory mechanism (*45, 55*). Through differential binding to three operator sequences (O_R_1, O_R_2, O_R_3), CI maintains lysogeny by binding preferentially to O_R_1 (adjacent to the lytic operon), while Cro promotes lysis by preferentially binding to O_R_3 (adjacent to the lysogenic operon). We show that Cro_Φ1720a-02_ mimics the CI role in the ϕNP prophage, by binding the operator adjacent to the lytic operon, repressing its transcription. This is a particularly elegant strategy employed by the ϕ1720a-02 phage, as it repurposes this fundamental regulatory mechanism to execute opposing effects on competing prophages.

Furthermore, we observe that Cro_Φ1720a-02_ constitutively inhibits ϕNP lytic operon expression even in the absence of an inducing trigger. Thus, upon activation of the SOS response, the ϕNP cannot be induced because Cro_Φ1720a-02_, which is not cleaved by LexA, is already bound to ϕNP lytic operator. This aligns with the reports that Stx prophages generally exhibit a higher rate of basal induction compared to phage lambda and other lambdoid phages (50). As a result, ϕ1720a-02 phage secures a competitive advantage by preventing the competing prophage from transitioning to the lytic cycle upon induction.

We found that Cro_Φ1720a-02_ is highly conserved among other Stx phages, suggesting that this repressor might exert its dual function across various host strains encountered by these phages. This raises the possibility that other Cro repressors may also regulate multiple elements, whether bacterial or within prophages, a prospect that requires further investigations.

Previously described prophage-prophage interactions involve small anti-repressor proteins that specifically bind to non-cognate repressors and trigger induction of co-hosted prophages (*56*–*58*). Other prophage cross-regulation mechanisms target general bacterial pathways like the SOS response or the translation machinery, making them nonspecific to individual prophages or elements (*53, 59*–*61*). Here, we show that Cro_Φ1720a-02_ directly targets a phage with a specific operator sequence in its control region, demonstrating a targeted cross-regulatory function. However, lysogenization of various *E. coli* strains with ϕ1720a-02 altered the induction profile of multiple different prophages within each lysogen tested. This lysogenization resulted in diverse effects on co-hosted prophages, ranging from inhibition to enhanced induction at different levels. This indicates that ϕ1720a-02 uses multiple approaches to interfere with the activity of co-hosted elements, by pathways that remain to be uncovered.

Our work implies that prophage cross-regulation is integral to the propagation and transmission of Stx prophages. We propose that prophage-prophage interference contributes to the evolution and ecology of Stx bacteria and plays a role in the emergence of new pathogenic strains.

## Acknowledgment

We thank Dr. Netanel Tzarum for his help and guidance with the structural analysis. Work in the Litvak laboratory is supported by Israel Science Foundation grant 2350/20, Ministry of Innovation, Science & Technology awards 3-17966 and 0005996, and by award 2019136 from the U.S.-Israel Binational Science Foundation.

