## Supplementary figures and tables for "The bully phage: A Shiga toxin-encoding prophage interferes with the induction of co-hosted prophages"

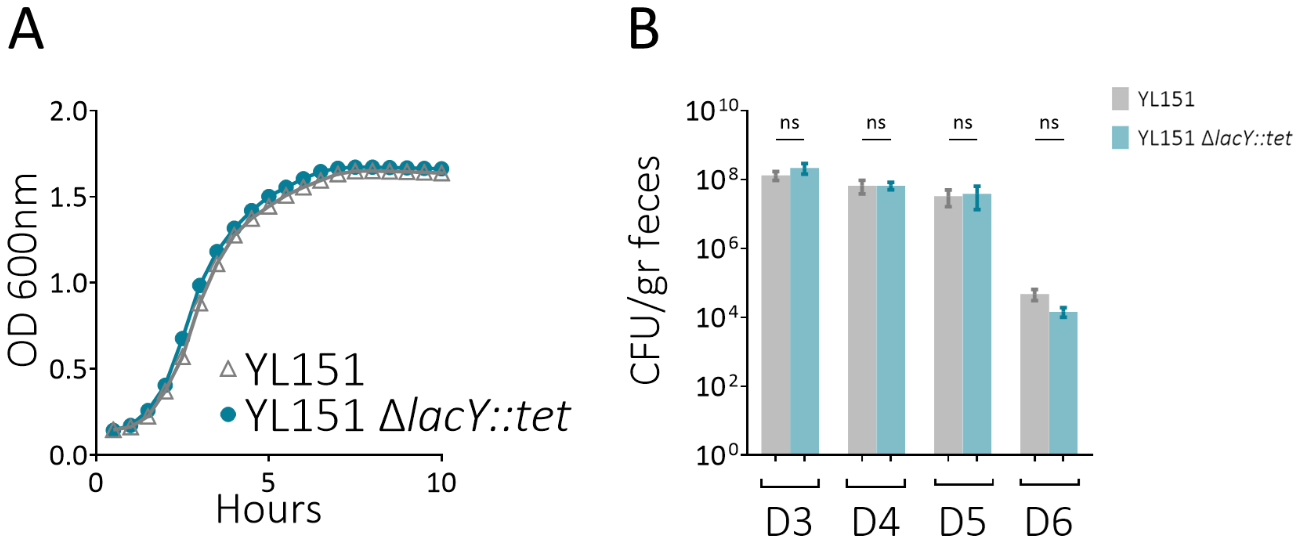


**Figure S1- Fitness of commensal *E. coli* and *lacY* mutant** (A) Growth curves of *E. coli* YL151 and *E. coli* YL151 Δ*lacY::tet* mutant in LB media measured by OD600nm. (B) Bacterial load in mouse feces following infection of mice with *E. coli* YL151 or *E. coli* YL151 Δ*lacY::tet* mutant , measured as CFU per gram of feces. Bars represent means ± standard error. ns=non-significant.

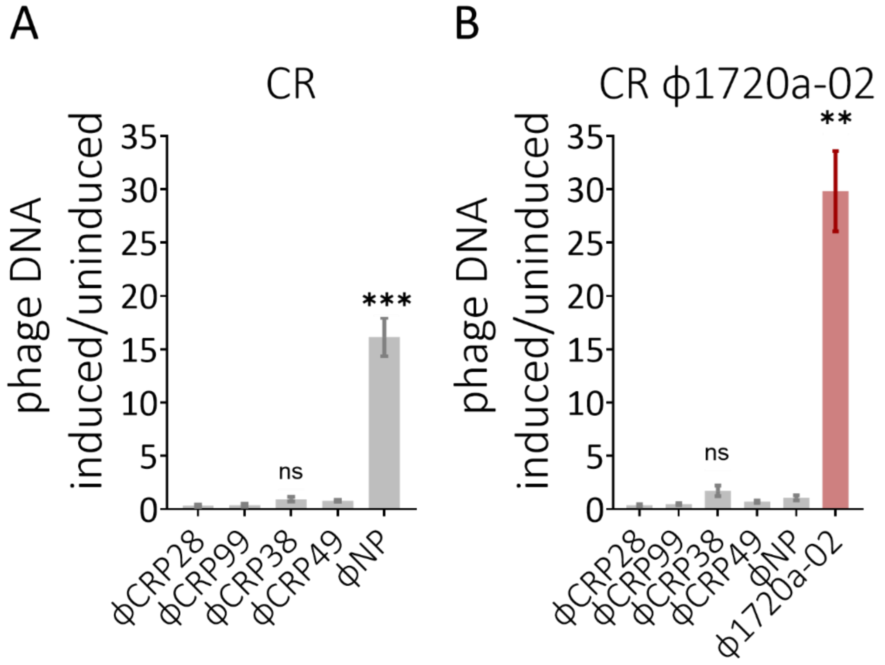


**Figure S2 - ϕNP induction by UV is inhibited when co-hosted with ϕ1720a-02 -** DNA levels of prophages encoded by (A) CR or (B) CR ϕ1720a-02 measured by qPCR. DNA levels are presented as relative copies in UV-treated vs. untreated cultures, normalized to bacterial DNA. Bars represent means ± standard error. *P<0.05. **P<0.002. ***P<0.001. ns=non-significant.

| | Strain | Identifier | Sequence |
|---|---|---|---|
| 1 | **ϕ1720a-02** | **KF030445.1** | **MQSPLRNVRKAHGFTLQHVAAGVQVNPATLSRIERLEQIPSIDLAERLANFFKGEISEMQILYPARFQSSQNRNELKPQEKEVSRG** |
| 2 | STEC316 | CP041431.1 | MQSPLRNVRKAHGFTLQHVAAGVQVNPATLSRIERLEQIPSIDLAERLANFFKGEISEMQILYPARFQSSQNQNGFKPQEQEVSRG |
| 3 | STEC409 | CP041422.1 | MQSPLRNVRKAHGFTLQHVAAGVQVNPATLSRIERLEQIPSIDLAERLANFFKGEISEMQILYPARFQSSQNQNGFKPQEQEVSRG |
| 4 | STEC435 | CP129348.1 | MQSPLRNVRKAHGFTLQHVAAGVQVNPATLSRIERLEQIPSIDLAERLANFFKGEISEMQILYPARFQSSQNQNGFKPQEQEVSRG |
| 5 | O170:H18 | CP076706.1 | MQSPLRNVRKAHGFTLQHVAAGVQVNPATLSRIERLEQIPSIDLAERLANFFKGEISEMQILYPARFQSSQNQNGFKPQEQEVSRG |
| 6 | STEC1155 | CP101978.1 | MQSPLRNVRKAHGFTLQHVAAGVQVNPATLSRIERLEQIPSIDLAERLANFFKGEISEMQILYPARFQSSQNQNGFKPQEQEVSRG |
| 7 | O157:H7 | AP018488.1 | MQSPLRNVRKAHGFTLQHVAAGVQVNPATLSRIERLEQIPSIDLAERLANFFKGEISEMQILYPARFQSSQNRNELKPQEKEVSRG |
| 8 | O22:H8 | CP023165.1 | MQSPLRNVRKAHGFTLQHVAAGVQVNPATLSRIERLEQIPSIDLAERLANFFKGEISEMQILYPARFQSSQNRNELKPQEKEVSRG |
| 9 | STEC1588 | CP129263.1 | MQSPLRNVRKAHGFTLQHVAAGVQVNPATLSRIERLEQIPSIDLAERLANFFKGEISEMQILYPARFQSSQNRNELKPQEKEVSRG |
| 10 | STEC2018-553 | CP075665.1 | MQSPLRNVRKAHGFTLQHVAAGVQVNPATLSRIERLEQIPSIDLAERLANFFKGEISEMQILYPARFQSSQNRNELKPQEKEVSRG |
| 11 | 2012EL | CP027586.1 | MQSPLRNVRKAHGFTLQHVAAGVQVNPATLSRIERLEQIPSIDLAERLANFFKGEISEMQILYPARFQSSQNRNELKPQEKEVSRG |
| 12 | O91:H21 | CP031906.1 | MQSPLRNVRKAHGFTLQHVAAGVQVNPATLSRIERLEQIPSIDLAERLANFFKGEISEMQILYPARFQSSQNRNELKPQEKEVSRG |
| 13 | 89-3506 | CP027520.1 | MQSPLRNVRKAHGFTLQHVAAGVQVNPATLSRIERLEQIPSIDLAERLANFFKGEISEMQILYPARFQSSQNRNELKPQEKEVSRG |
| 14 | O178:H19 | CP024289.1 | MQSPLRNVRKAHGFTLQHVAAGVQVNPATLSRIERLEQIPSIDLAERLANFFKGEISEMQILYPARFQSSQNRNELKPQEKEVSRG |
| 15 | EH41 | CP045213.1 | MQSPLRNVRKSHGFTLQHVAAGVQVNPATLSRIERLEQIPSIDLAERLANFFKGEISEMQILYPARFQSSQNRNELKPQEKEVSRG |
| 16 | JNE141411 | AP027256.1 | MQSPLRNVRKSHGFTLQHVAAGVQVNPATLSRIERLEQIPSIDLAERLANFFKGEISEMQILYPARFQSSQNRNELKPQEKEVSRG |
| 17 | O113:H21 | CP031892.1 | MQSPLRNVRKSHGFTLQHVAAGVQVNPATLSRIERLEQIPSIDLAERLANFFKGEISEMQILYPARFQSSQNRNELKPQEKEVSRG |

**Figure S3- Homologs of Croϕ1720a-02 in other Stx phages -** Alignment of Cro protein sequences from Stx phage-carrying bacteria. The conserved Arginine at position 32 is highlighted in blue, non-conserved amino acids are shaded in red.

**Supplementary Tables**

**Table S1. Bacterial strains used in this study**

| | | |
|---|---|---|
| ***C. rodentium*** | | |
| DBS100 | *Wild type* | ATCC 51459 |
| DBS770 | *C. rodentium* ϕ1720a-02-*cat* | ATCC BAA-2623 |
| TW480 | DBS770 Δ*stx::kan* | This study |
| RU355 | DBS770 Δ*lyt::kan* | This study |
| RU538 | DBS770 Δ*cro::kan* | This study |
| ***E. coli*** | | |
| DH5α | *fhuA2 lac(del)U169 phoA glnV44 Φ80' lacZ(del)M15 gyrA96 recA1 relA1 endA1 thi-1 hsdR17* | Lab collection |
| MFD λ pir | MG1655 *RP4-2-Tc::[ΔMu1::aac(3)IV-ΔaphA-Δnic35-ΔMu2::zeo] ΔdapA::(erm-pir) ΔrecA.* | (*62*) |
| YL151 | Wild type natural isolate from mouse feces | This study |
| YL262 | Wild type natural isolate from mouse feces | This study |
| YL256 | Wild type natural isolate from mouse feces | This study |
| TW505 | YL151 Δ*lacY::tet* | This study |
| TW459 | YL151 ϕ1720a-02-*cat/pCAL61* | This study |
| TW482 | YL262 Δ*lacY::tet/pCAL62* | This study |
| TW499 | YL262 Δ*lacY::tet* ϕ1720a-02-*cat/pCAL62* | This study |
| TW485 | YL256 Δ*lacY::tet/pCAL62* | This study |
| TW502 | YL256 Δ*lacY::tet ϕ1720a-02-cat /pCAL62* | This study |

**Table S2. Plasmids used in this study**

| **Plasmid** | **Identifier** | **Description** | **Reference** |
|---|---|---|---|
| pKD46-strep | pKD46 | λ Red recombinase expression | (*46*) |
| pCAL61 | pCAL61 | pWSK129 carrying the omega cassette | (*63*) |
| pEP185.*2* | pEP185.*2* | *ori*R6K *mobRP4 cat* | (*64*) |
| pCAL62 | pCAL62 | pWSK29 carrying the omega cassette | (*63*) |
| pUC19 | pUC19 | pUC6 *lacZA ampR* | (*65*) |
| pUC19-*stx::kan* | pTW475 | pUC19 with *stxAB* homologous regions flanking a *kan* cassette | This study |
| pBAD-*hisD*-TagRFP675 | pRU514 | pBAD/HisD *araC ampR TagRFP675* | Addgene #44274 |
| pBAD-*cro* $_{\phi 1720a\text{-}02}$ | pRU517 | pBAD-*hisD*-TagRFP675 in which the TagRFP675 was replaced with His6 tag and *cro* $_{\phi 1720a\text{-}02}$ | This study |
| pEP185.2-*cro::kan* | pRU533 | pEP185.2 with *cro* homologous regions flanking a *kan* cassette | This study |
| *pBR-C5-5-luxPl* | pRU565 | *pBR322* containing the *Photorhabdus luminescens luxCDABE* genes | (*36*) |
| p*C* $_{\phi NP}$-*lux* | pRU575 | *pBR-C55-luxPl* in which the C5-5 promoter was replaced with the ϕNP lytic control region | This study |
| pACYC184 | pRU653 | *p15A cat tet* | ATCC #37033 |
| pACYC184-*pC* $_{\phi NP}$-*lux* | pRU654 | The pACYC184 backbone with the *luxPl* operon downstream to the ϕNP lytic operator | This study |

**Table S3. Primers used in this study**

| Identifier | Name | Sequence (5'→3') |
|---|---|---|
| 323 | *lacY::tet* F | gcgatcattccgccctgatatgttggtcggataaggcgctcgcgccgcatccgacattgattgcttaagcgacttcattcacctgacgacgcagtagaagacccactttcacatttaag |
| 324 | *lacY::tet* R | ggagcccgtcagtatcggcggaattccagctgagcgccggtcgctaccattaccagttggtctggtgtcaaaaataataataaccgggcaggctatgtctgcccgtatttcgcgtaaggaaatccattatgtactatttaaaaaacacaaacttttggctaagcacttgtctcctgttta |
| 403 | *lyt::kan* R | attacttcagccaaaaggaacacctgtatatgaagtgtatattatttaaatgggtactgtagagcgcttttgaagctggg |
| 404 | *lyt::kan* F | gtcgggctgcggtctctgttaatgagggaatacagcgacgatacggcgcatcagcaaaacttgatcccctcacgctgccg |
| 254 | *kan* F | gatcccctcacgctgccg |
| 255 | *kan* R | agagcgcttttgaagctggg |
| 485 | pUC19 F | ggtgcctcactgattaagcattggtaac |
| 486 | pUC19 R | caaataggggttccgcgcac |
| 738 | P1 *stx* overlap pUC19 F | gcttaatcagtgaggcaccgctttcggatggttaagg |
| 739 | P2 *stx* overlap *kan* R | cggcagcgtgaggggatcgacggtaacaatcaaatc |
| 740 | P3 *stx* overlap *kan* F | cagcttcaaaagcgctctggtcgaaaagtctatcgtaaactccc |
| 741 | P4 *stx* overlap pUC19 R | gcgcggaacccctatttggatgccaggtatgaggcgaag |
| 643 | pBAD33 R | atggtgatggtgatggtggc |
| 644 | pBAD33 F | ccagcttcatgtacggcag |
| 835 | pBAD33 RBS x6HIS R | agatctgccatggtgatggtgatggtggctccccatggttaattcctcctgctagcccaaaaaaacggg |
| 836 | pBAD33 F | ggctgttttggcggatgag |
| 837 | *Cro* overlap x6HIS F | ctctcatccgccaaaacagccttacccacggcttacctc |
| 838 | *Cro* overlap pBAD33 R | catcaccatggcagatctgtgggtaagcatcactgg |
| 513 | *kan* David F | ttctgctccctcgctcaggggggaaagccacgttgtgtc |
| 514 | *kan* David R | cagggagcactggtcaacccagccagaaagtgagggag |
| 578 | pEP185.2 F | ccactagttctagagcgg |
| 579 | pEP185.2 R | gataccgtcgacctcgag |
| 869 | P1 *cro* overlap pEP185.2 F | ctcgaggtcgacggtatcggttaattgtgtcgcttaagg |
| 870 | P2 *cro* overlap David R | gttgaccagtgctccctgataacctccacatattgcgtg |
| 871 | P3 *cro* overlap David F | ctgagcgagggagcagaagcatcactggaaagtggaaaaac |
| 872 | P4 *cro* overlap pEP185.2 R | ccgctctagaactagtggcagtcaggtaaagttctctg |
| 885 | pBRlux overlap pCϕNP *lyt* F | tacttacgacttgcgcaagttgctttgcatatcacgcaaaaacaacctgtaatagaacgcgtacccggggatccc |
| 886 | pBRlux overlap pCϕNP *lyt* R | ttgcgcaagtcgtaagtacacattaatatgcgtattgcgcaattatagggggctttcaggaggggcaaatatgac |
| 894 | *lux* R | ccagtaagtaattacttgagtcgac |
| 939 | *lux* F | ctatagggcgaattcctttaa |

| | | |
|---|---|---|
| 940 | pACYC184 overlap *lux* F | gaattcgccctatagggcagttattggtgccctt |
| 941 | pACYC184 overlap *lux* R | gtaattacttactggcagggcttcccggtatcaac |
| 41 | *eae* F | gga agc caa agc gca caa |
| 42 | *eae* R | ggc gcg agc ggt cac ttt ata aac |
| 47 | *StxA2* F | agttctgcgttttgtcactgtc |
| 48 | *StxA2* R | cggaagcacattgctgatt |
| 81 | ϕStx *attP* F | cacagtgttttgcatctcacac |
| 82 | ϕStx *attP* R | acctaacgcgagaaaaatagcc |
| 677 | ϕNP F | caatttccaccccagagcag |
| 678 | ϕNP R | tctgctgttgggctatgact |
| 786 | CR *RpoA* F | acgtcagccggaagtgaaagaaga |
| 787 | CR *RpoA* R | agcggacagtcaattccagatcgt |
| 857 | ϕNP *attP* F | gctatgagctagacgactac |
| 858 | ϕNP *attP* R | ctgagaggctggcttgattg |
| 935 | ϕNP *holin* F | aattagcagggaggccgaag |
| 936 | ϕNP *holin* R | atcgcgatgtacctcccttc |
| 502 | ϕSM F | gcagcttcagtgataaggcc |
| 503 | ϕSM R | gtccgcctttatcgaaccac |
| 975 | ϕSM *attP* F | cagtattgcccgtctaaatggtt |
| 976 | ϕSM *attP* R | ctgtcaaacgcgctaaaacc |
| 498 | ϕCrp28 F | gcccgctctgttaatgtctg |
| 499 | ϕCrp28 R | aaacgcgaggctaacgaatc |
| 500 | ϕCrp99F | gaatcagtaacagcaggccg |
| 501 | ϕCrp99 R | actgaattccgctggtacca |
| 504 | ϕCrp49 F | tcgctacgatttgccgaaag |
| 505 | ϕCrp49 R | tcttctggtgcattccccat |
